# P3ANUT: An enhanced DNA sequencing analysis platform for uncovering and correcting errors in peptide display library screening

**DOI:** 10.1101/2025.05.12.648809

**Authors:** Liam Tucker, Ethan Koland, Jasmyn Gooding, Hassan Boudjelal, Sanaz Ahmadipour, Robert A. Field, Derek T Warren, David Baker, Yingliang Ma, Maria J Marin, Taoyang Wu, Chris J Morris

## Abstract

Display technologies are used extensively in the discovery of peptides and antibodies towards the development of new medicines and diagnostic tools. Phage display technology enables the filtering of <10^10^ random peptides/antibodies down to and enriched pool of candidates with favourable binding affinities. In recent years, next-generation DNA sequencing technologies have increased the precision and accuracy with which the peptide sequences of phage display library clones are identified. Inaccuracies in DNA sequencing such as substitutions, insertions and deletions in the library oligonucleotide region have the potential to result in the identification of erroneous candidate sequences. Here, we describe a <u>P</u>ython <u>P</u>ipeline for <u>P</u>hage <u>A</u>nalysis through a <u>N</u>ormative <u>U</u>nified <u>T</u>oolset (P3ANUT) which employs Levenshtein distance, k-mer approaches and a novel encoding scheme on paired-end sequencing outputs to correct sequencing errors from next-generation sequencing outputs of display library screens. We introduce an easy-to-use and highly customisable computational tool with graphical user- and command line interfaces to process entire datasets within a single input, as well as visualisation tools for candidate analysis and data generation. P3ANUT shows significant improvements in read recovery, overall read quality, and runtime compared to a previously published pipeline.

**GRAPHICAL ABSTRACT:** 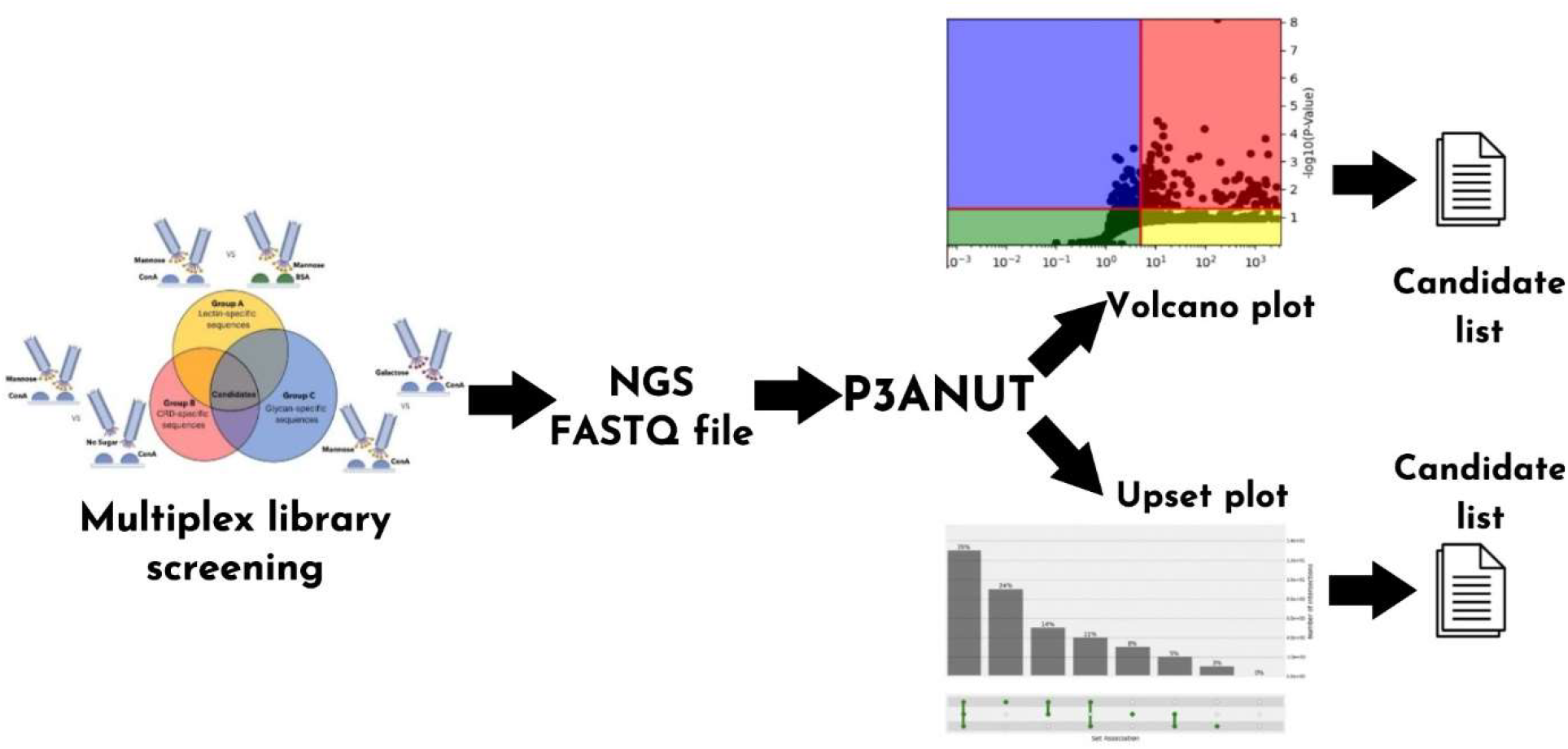

## 1. INTRODUCTION

Display technologies is a term that encompasses multiple techniques, such as phage-, ribosome-, and yeast display. They are recombinant screening techniques that afford the screening of large libraries of peptides and proteins to identify binding motifs for a specific target. Examples of such targets can span the length and complexity scales from small molecules to whole cell or tissue surfaces. The phage display approach involves the ectopic expression of peptides or protein fragment libraries on the coat proteins of bacteriophage virions (1,2). Peptide phage display typically involves 2-5 rounds of affinity selection against a given target in a process called “library panning”. After elution of binding clones from the target they undergo clonal amplification in a suitable bacterial host, before conducting phage DNA extraction and sequencing of the coat protein gene region that includes the randomised recombinant fragment. From this, the peptide sequences that mediate target engagement are deduced. Discovery of binding peptides can provide lead candidates for drug development as exemplified by the approved kallikrein inhibitor drug, ecallantide, which was FDA approved in 2009 for the treatment of hereditary angioedema. In oncology both antibody and peptide phage display has been utilised to discover potential new treatments highlighting the diversity of applications display technologies presents.

Since its invention in the 1980s, peptide phage display technology has developed significantly in terms of both the number of randomised peptides (i.e. library complexity) as well as through increased chemical diversity. Changes in the chemical diversity of randomised native peptides has been achieved in several ways. These include the site-specific attachment of other chemical species such as reactive glycans to the N-terminus. Complex cyclic peptide topologies including bi- and tri-cyclic motifs have been achieved either through the chemical reactivity of cysteine with, for example, tris(bromomethyl) benzene to form bicyclic libraries, or directed folding through the inclusion of peptide sequences (e.g. CPPC) that promote the formation of a cystine-knot topology (3). These increases in library complexity and the increased pressure for high-throughput screens against drug targets has demanded faster and more accurate DNA sequencing to ensure the correct candidate sequences are pursued.

High-throughput sequencing (HTS) technologies have massively increased the depth and coverage of phage display library sequences that emerge from any given panning experiment. HTS of peptide phage display outputs can produce extremely large volumes of DNA sequencing data (approximately 600 MB per dataset). After processing these data can reveal >1,000 peptide sequences of varying abundances after a single library panning round. This has significantly increased the power and speed of recombinant display library screens. Concomitant with an increased surveillance of phage library diversity across consecutive panning rounds is the potential for distraction by either non-specific binding clones or phage clones that display a marginal growth advantage over other library clones, so called parasitic sequences (4). Non-specific binding clones can be selected against by protocol modification to enrich for specific binders. However, parasitic clones are an innate element of phage display libraries due to the randomised nature of the recombinant DNA included in their cloning.

The key aim of HTS of phage display outputs is to identify the abundance of specific sequences rather than consensus generation. Therefore, sequencing errors can have a profound effect on results because read length is short and linking of non-contiguous sequences, a process known as scaffolding, is not possible. Earlier MatLab-based software developed by Rebollo et al. (5), Matochko et al. (4) and He et al. (6) processes phage display HTS data and undertakes sequence clustering and candidate filtration. Each of these scripts has their own distinct features, yet there remains a need to deal with sequencing errors to improve sequence assembly confidence. Such errors include substitution errors that are caused by noise in the fluorescence signals generated during sequencing, which interferes with the base-calling process. This can either be a silent substitution where the base substitution translates into the same amino acid, or otherwise generates an entirely different amino acid when translated. More serious errors such as insertions and deletions (otherwise known as indels) can arise rarely due to failure to fully remove free bases in solution or remove fluorescent terminators from annealed bases, causing the formation of a leading/lagging signal, respectively (7). Indels in sequencing will cause frameshifts that result in the amino acids generated ahead of the indel base to be entirely misread. The unfeasibility of sequence scaffolding means that alternative methods are needed to mitigate the impact of these sequencing errors on the results generated from phage display HTS.

HTS has seen many developments over time to reduce the natural analogue noise found in sequencing, with most of these developments found in genomic sequencing. As these problems can be thought of as string problems, techniques applied to such can also be applied to sequence processing in the form of edit distance, in which pairwise alignments of two strings details the number of steps required to make both strings match. One such approach utilised is Levenshtein distance (8), a metric of edit distance that considers substitutions, insertions and deletions, in turn providing a simple algorithmic approach to process all errors that can be detected in peptide display HTS (9). Essentially, it reports the number of substitutions, insertions and deletions necessary to match two different DNA sequences. For example, there is a Levenshtein distance of 2 between the terms, MOP and MAD, since O must change to A and P must change to D. The Levenshtein distance approach has been successfully deployed in sequence alignment and as a foundation for later, more extensive developments such as the Smith-Waterman algorithm used by BLAST (10). In the HTS analysis of phage display output, the simplicity of Levenshtein distance pairwise analysis on the forward and reverse reads is an attractive approach to increase sequencing precision and increased accuracy in the identification of candidate peptides for further development.

While Levenshtein distance provides the means of detecting disagreements in pairwise matching, either sequence can contribute to the construction of the final “consensus” sequence. Thus, a means of determining the most likely correct base in a disagreement is necessary. Quality scores, generated as part of the FASTQ format, provide a confidence to the called base being correct from the analogue fluorescence signals generated during sequencing, in turn resulting in scores showing fuzziness. Indels can be resolved in peptide display HTS with little issue because the expected length and general library composition is known. However, this cannot be used for substitutions occurring in the variable region of the library. In cases where a base disagreement shows very little or no difference in quality scores between the bases, a decision involving only the scores directly involved can lead to error-prone corrections, due to the fuzziness described above (11). To overcome this issue, group comparisons involving nearby bases as *k*-mers are utilised. This compares each k-mer group containing the base disagreement, with the base chosen being from the side with the most *k*-mer groups with greater quality scores against their comparison. Such an approach has been deployed successfully in genomic sequencing with CASPER (12). Compared with other tools for the same purpose CASPER displays greatly improved accuracy and F_1_ score metrics in tested datasets, proving the utility of this approach. Here, we hypothesised that a combined approach of Levenshtein distance matching and k-mer group comparisons could be employed in peptide display sequencing analysis to improve sequencing fidelity and overall performance of the sequencing process. Compared with other tools for the same purpose CASPER displays greatly improved accuracy and F_1_ score metrics in tested datasets, proving the utility of this approach.

In this work, we have developed the <u>P</u>ython <u>P</u>ipeline for <u>P</u>hage <u>A</u>nalysis through a <u>N</u>ormative <u>U</u>nified <u>T</u>oolset (P3ANUT) software which processes sequencing data with a high level of data quality while maintaining runtimes comparable to or better than the previously published pipeline comparators. The software was used to analyse the outputs from an exemplar post-translationally modified glycopeptide phage display screen against a glycan binding lectin, concanavalin A (ConA). The inclusion of analytical tools as part of the software enables the filtration of undesirable peptides (non-specific binders, etc) and the generation of LOGO sequence data integrates well with existing tools (e.g. Clustal Omega) to discover consensus motifs and candidate sequences for further development.

## 2. MATERIAL AND METHODS Reagents & Biological Resources

The mannose-binding lectin, concanavalin A (ConA) from Canivalia ensiformis was from Vector Laboratories. Bovine serum albumin (BSA) was purchased from Fisher. Synthetic library oligonucleotides were purchased from Sigma-Aldrich. Restriction endonucleases, EagI-HF (R3505) and KpnI-HF (R3142), Klenow fragment (M0210), T4 DNA ligase (M0202), NEBuffer 2 (B7002), rCutSmart Buffer (B6004), and M13KE phage (E8101) were purchased from NEB. Platinum™ SuperFi II DNA Polymerase (12361010) and SuperFi™ Buffer (12355005) were purchased from ThermoFisher Scientific. 96-well Microplates were purchased from Corning (3370). All other reagents were of analytical grade quality and from Sigma-Aldrich or Fisher Scientific.

### Library Design and Construction

A disulfide constrained, 7-mer library was prepared using standard cloning methods as described in the product manual for the NEB Ph.D Phage Display Cloning System (E8101).

The library oligonucleotide was:

5’-CCCGGGTACCTTTCTATTCTCACTCTTCTTGT(NNK)_7_TGTGGTGGAGGTTCGGCCGGGCGC-3’

Which translates to SC(X)_7_CGGG, which includes an N terminal serine residue after the peptidase leader sequence which enables the oxidation of the N terminus and easy attachment of reactive mannose glycan through oxime ligation (13).

Libraries were annealed with 3 molar equivalents of universal extension primer in 50 µL TE buffer with 100 mM NaCl by heating to 95 °C for 2 minutes, then cooled to 37 °C over 15 minutes. The annealed library was extended in NEBuffer 2 with 10 mM dNTPs and Klenow fragment (NEB #M0210), and double digested in NEB CutSmart Buffer using EagI-HF and KpnI-HF. The library was then verified for complete digestion and purified using 9% PAGE gel purification according to procedures listed in the NEB Ph.D. Phage Display manual. The purified digested libraries were mixed with digested and agarose gel purified M13KE phage genome and ligated using T4 DNA ligase. The ligated vector was then added to freshly thawed electrocompetent ER2738 E. coli cells and transformation was carried out on a BioRad Gene Pulser at 25 µF, 200 Ω, and a maximum of 2.5 kV then recovered all in accordance with the procedures listed in the NEB Ph.D. Phage Display manual.

### Library Panning

The SCX_7_C library was glycosylated at the N terminus by the attachment of either aminooxy-galactose or aminooxy-mannose via oxime ligation using the protocol described in Ng et. al. (13) with minor alterations that are described in the Supplementary Methods. Each of the proteins listed in Table was non-specifically adsorbed to three wells of a polystyrene 96-well plate, at the concentrations specified for each of the panning rounds.

**Table 1.** Phage library panning targets

| Target | Origin | Sugar binding preference | Ions required | Concentration ($\mu\text{g.mL}^{-1}$) | |
| --- | --- | --- | --- | --- | --- |
|  |  |  |  | Round 1 | Round 2 |
| **ConA** | *Canivalia ensiformis* lectin | Mannose | $\text{Ca}^{2+}$, $\text{Mn}^{2+}$ | 50 | 25 |
| **BSA** | Bovine serum albumin | N/A | N/A | 50 | 25 |

Library panning was performed against each target protein as described in detail in the Supplementary Methods. Washes were performed using TBS + 0.1% Tween-20 for six washes in round one and TBS + 0.5% Tween-20 for six washes in round two. Between rounds, 75 µL of phage eluate was taken forward for amplification while 30 µL was subjected to DNA extraction to produce the Round 1 unamplified phage datasets. The entire volume of Round 2 eluates were subjected to DNA extraction and DNA sequencing as described in the Supplementary Methods.

## 3. RESULTS

### General pipeline workflow

Phage display HTS data, generated using Illumina technology, were processed using a collection of Python scripts supported by a user interface for improved ease of use and compiled as P3ANUT (https://github.com/taoyangwu/P3ANUT). The overall pipeline structure is shown in Figure 1. P3ANUT comprises five key modules that assemble and decode DNA sequences to compile peptide sequence lists. The amino acid data was then summarized to a descending order list of peptides sorted by abundance. Sequencing data from different experimental groups and technical replicates were combined to provide means and standard deviations and analysed by t-test to filter out significantly enriched phage clones presented as a combination of volcano and UpSet plots.

**Figure 1.**
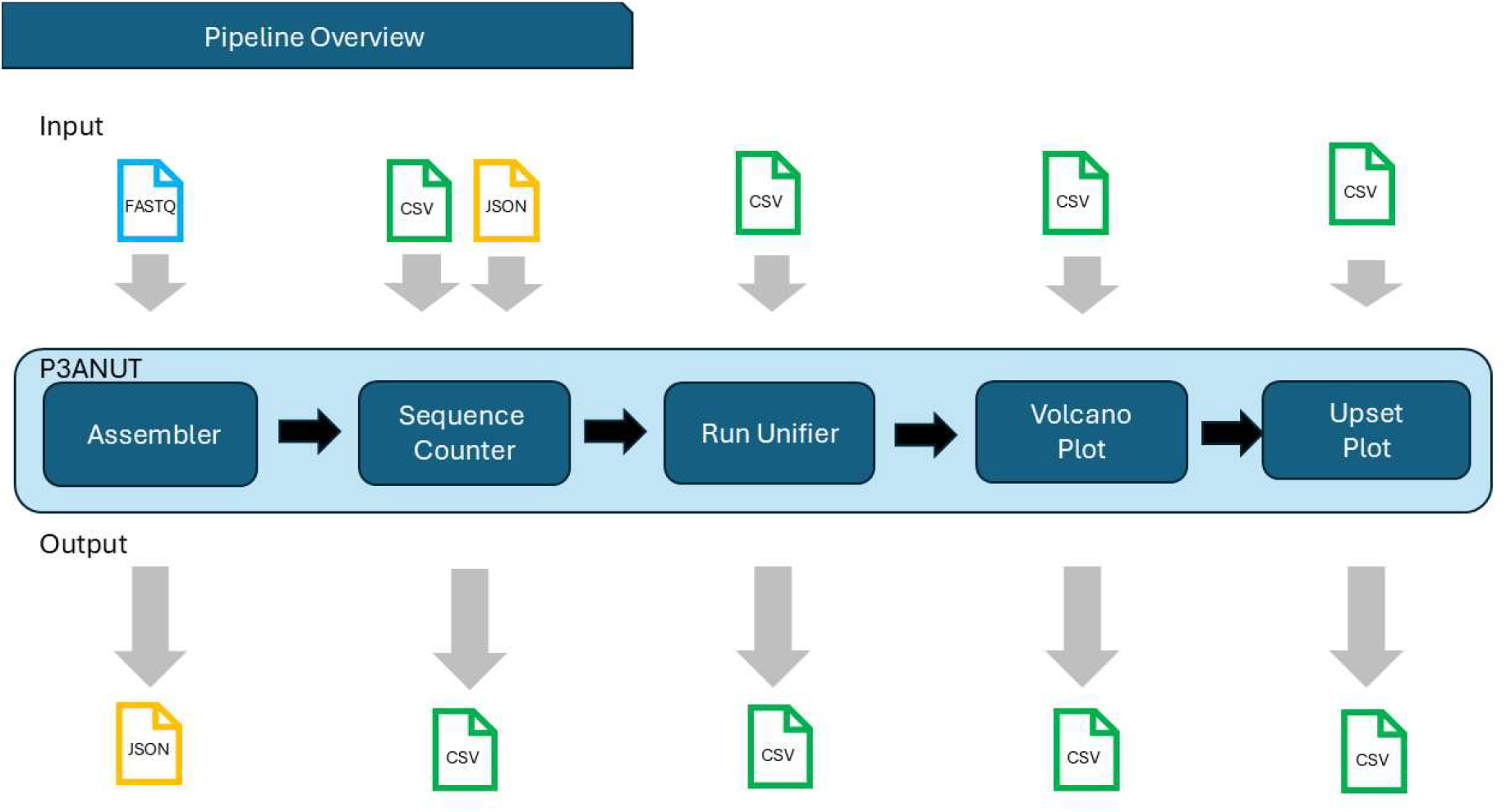
The Overall P3ANUT Pipeline. The overall pipeline of P3ANUT is made of five primary components for general use. From the Sequence Counter onwards, a general data type is used that can be used at any point onwards, if necessary. The assembler processes paired forward and reverse sequencing reads, encoding sequence and quality scores to reduce memory usage while efficiently merging and resolving mismatches to generate consensus sequences. Low-quality sequences are filtered, and consensus DNA is converted into amino acids using optimised algorithms for rapid and accurate downstream analysis. The sequence counter script processes DNA or amino acid datasets by filtering sequences based on user-defined criteria, such as counts or regex matches, and applying encoding and clustering methods to generate detailed outputs.

### Assembler

The NGS process typically generates two datasets as FASTQ files. One represents the 5’-3’ (forward) reads, while the other captures the 3’-5’ (reverse) reads. The reverse reads are converted to their reverse complement to facilitate pairing in the analysis pipeline. In cases where the output sequencing files do not include a reverse complement file, a simulated reverse file is produced using a small script that generates the reverse from the forward read. Subsequently, these datasets will not show errors, resulting in improved processing times due to no error correction processes taking place. Both datasets are filtered to retain sequences within a user-specified length range (e.g. 20-90 bp, according to the length of the library oligonucleotides), removing any trailing erroneous sequences, and subsequently trimmed to focus on the specified region of interest. Once paired, the sequences are then encoded using a novel overlapping 8-bit encoder (Figure 2 below and Algorithm 1 in Supplementary Information) to greatly reduce memory usage with little impact on runtime compared to existing methods (see Table 2).

**Figure 2.**
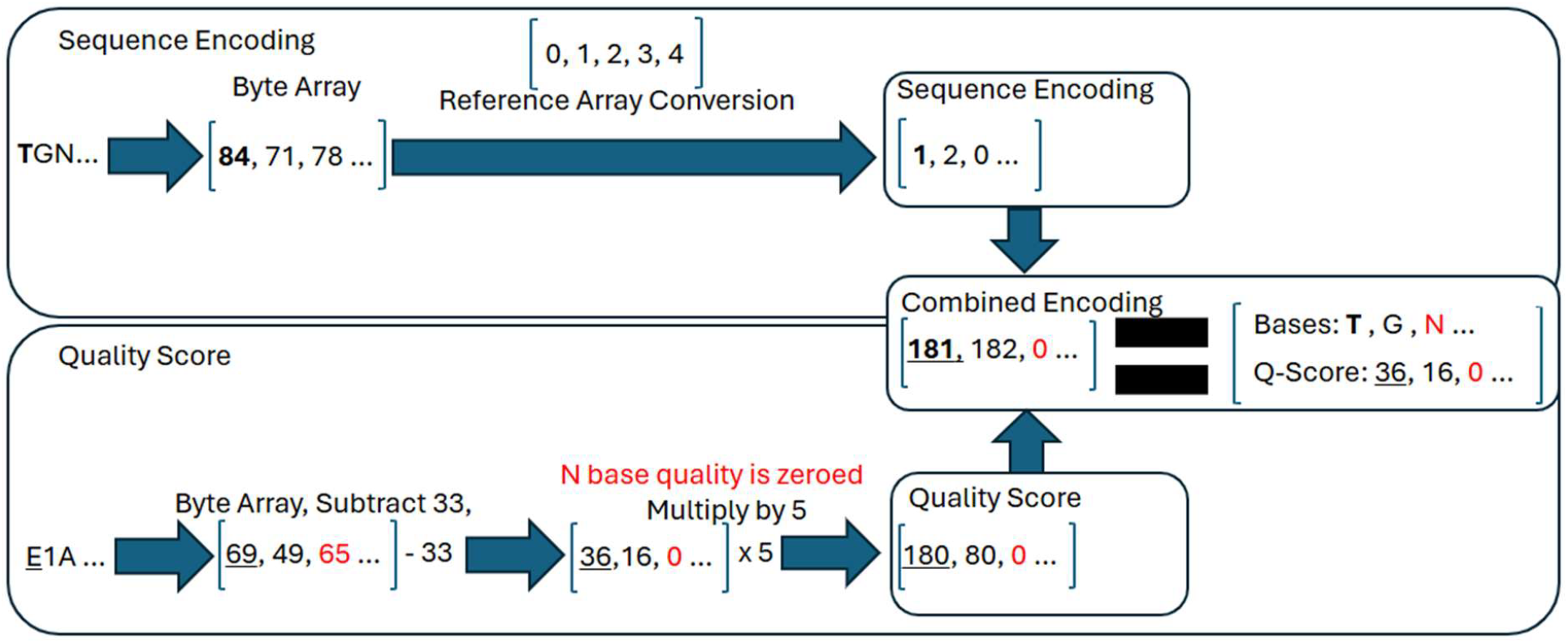
Example of Overlapping 8-Bit-Encoding in P3ANUT. First, each nucleotide base in the sequence is encoded by a number between 0 and 4. Secondly, the associated quality score is converted to a multiple of 5. Finally, these two numerical values are combined to create a unique representation of the pair. When the base is not ambiguous the resulting combined number is nonzero, which can be uniquely decode back to the pair. For example, the pair (T,E) highlighted in the figure is converted to the number, 181.

**Table 2.** Comparison of memory size and runtime for established Python coding methods compared to the new Overlapping 8 bit method. This compares the encoding time (in microseconds, µs) and merging time (i.e. time taken to merge) for four different methods.

| Method Name | Memory Size | Encoding Time ($\mu s$) | Merging Time ($\mu s$) |
| --- | --- | --- | --- |
| Python Char | 56 bytes | 49.9 | 15.6 |
| NumPy 8 Bit | 1 byte | 30.3 | 3.03 |
| NumPy 16 Bit | 2 bytes | 36.9 | 3.18 |
| Overlapping 8 Bit | 1 byte | 39.1 | 3.59 |

### DNA sequence analysis

For each of N (ambiguous base), T, G, C, A the index of the ASCII character is assigned the corresponding number 0, 1, 2, 3, 4. While maintaining this reference requires additional memory allocation during script compiling, it significantly reduces the runtime performance of sequence encoding. Another reference array is initialized for reverse-complement sequences. Maintaining a second reference array demands more memory, however it saves the need of an O(n) calculation to swap the bases with their matching base. Another implementation of the conversion involves the use of a polynomial. Each X (ASCII character codes) relates to a Y (encoded number). While both methods are O(c) run time, the polynomial involves significantly more complex and slower calculation.

N-bases are automatically reassigned a quality score of 0 to ensure these are encourage replacement with non-ambiguous bases. Thus, the consensus encoding will entail a value of 5Q + S, where Q is the quality score, and S is the respective base value. To decode the consensus scores, a similar operation of converting a string into an integer is performed. The integer representation of the ASCII characters needs to be converted to the Phred score, so all the scores are subtracted by 33. Maintaining 2 arrays to represent the sequences and their scores are memory intensive, through the memory overhead, and unused bits with the allocated bit. Python stores traditionally primitive objects, such as integers, in classes. The size of these default classes is 28 bytes but grows depending on the data size. Compressing and further encoding the sequences allows for reduced memory usage at small impact to runtime performance. NumPy arrays allow for the declaration of the bit allocated to storage of an integer.

The score and sequence can be encoded into a 16-bit integer. This represents a 97% reduction in the usage of memory per base-score pair. The first 3 bits are reserved for storing the encoded codon. 3 bits are needed to represent the 4 different possible bases and the N base. The remaining 13 bits are allocated to storing the sequence score. Encoding and decoding is performed quickly using bit-shift operations. Implementing a more complex encoding algorithm, the sequences can be further encoded into a single byte. The encoded Phred scores are multiplied by 5 and the encoded base sequences are added; resulting in the encoded consensus sequences. This operation can be easily decoded with a joint division, modulus operation. Depending on the chip architecture, this methodology takes 42 CPU cycles compared to bit segmentations that take 1 CPU cycle.

### Sequencing Error Correction

Figure 3 below shows the methods used to generate consensus sequences in phage display library sequencing results. More details can also be found in Algorithm 2-4 in Supplementary Information.

**Figure 3.**
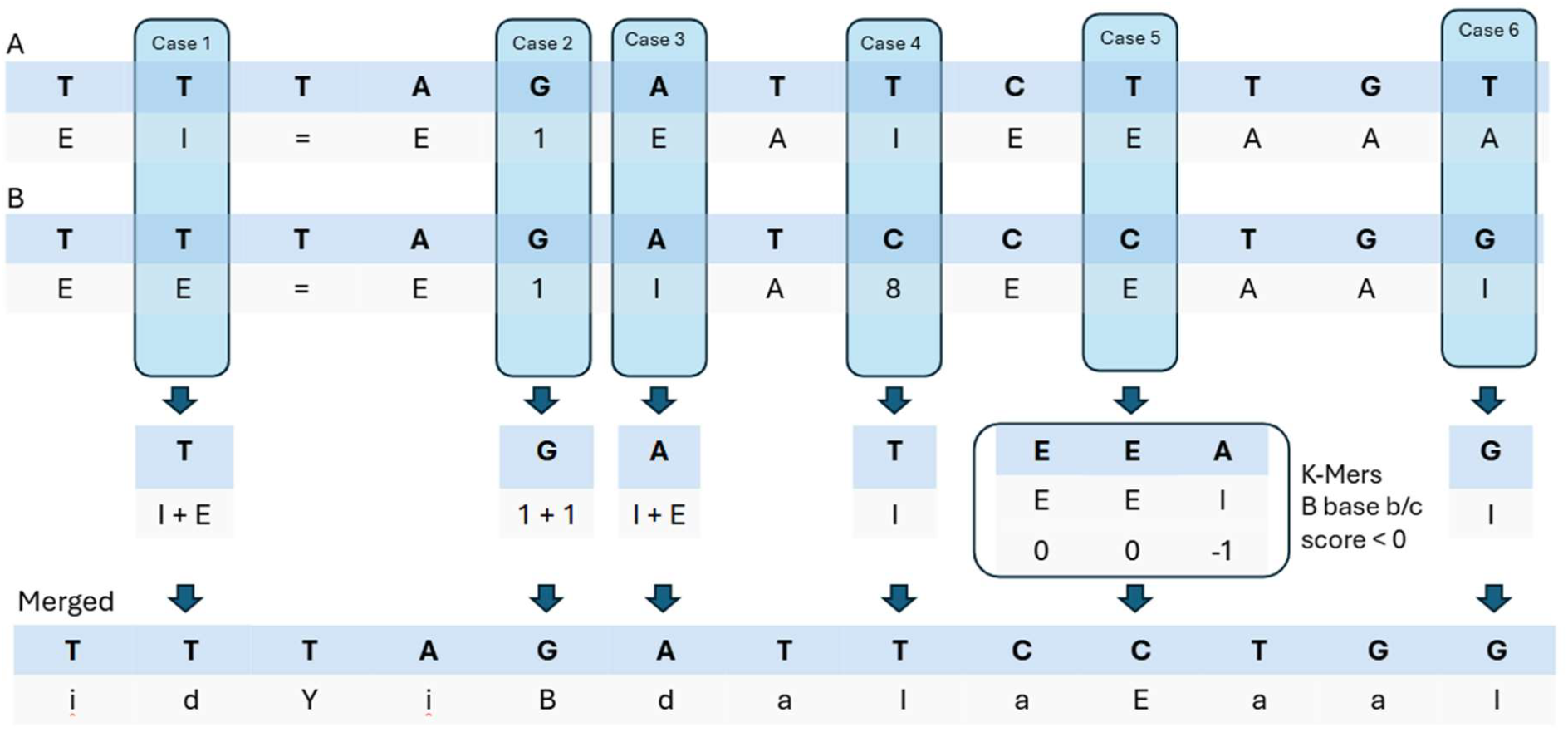
Example of Non-Indel Sequence Merging and Associated Cases in P3ANUT. When merging both forward (A) and reverse (B) reads to create a consensus (merged) sequence, various situations can arise from the presence of substitution errors which make up the majority of errors in HTS. Note that reads in B are first reversed and then complemented from the raw reverse read data. Cases 1 to 3 of above are carried out when compared bases match and in which case, the quality score is merged to represent the lowered chance of both reads being wrong simultaneously on the same base. Cases 4 and 6 represent substitution errors where the quality scores are above the difference threshold, in which case the base with the greater quality score is used for the merged sequence, assuming the quality score of that read. Case 5 represents a substitution error where the quality scores of the two bases are within the difference threshold, which are resolved by comparing the local group of bases as k-mers as first described by Kwon, Lee and Yoon (11).

In cases 1-3, matching base pairs have their quality scores combined under the following assumption:

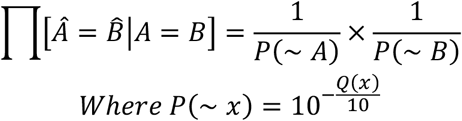

**Equation 1:** Approximated Incorrect Base Reporting Chance of a Matching Base-Pair

Where Â and 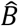 represent the true bases of the sequence and *A* and *B* represent the bases found through sequencing, *P*(*∼ A*) and *P*(*∼ B*) represent the error scores of the sequenced bases, and *Q*(*x*) represent the reported quality score. This means that if both bases had a quality score (*Q*(x)) of 30, which represents a 0.1% chance of the base being incorrect, the chance of both sequenced bases being incorrect is thus 0.0001%. Due to memory limits, the maximum cutoff for merged quality scores generated is 40 to maintain uniformity.

Where base disagreements arise, in cases of large quality differences, as seen in cases 4 and 6, the base with the greater quality score is chosen and this score is assumed in its place. In cases of non-significant or no quality score differences (Case 5), an alternative method is deployed, based on the group k-mer comparison methodology developed by Kwon et al (12). The comparison can be described as the following: Where base disagreements arise, in cases of large quality differences seen in cases 4 and 6, the base with the greater quality score is chosen and this score is assumed in its place. In cases of non-significant or no quality score differences an alternative method utilising group k-mer comparisons based on methodology seen in Kwon et al (12) as seen in case 5, where the comparison can be described as the following:

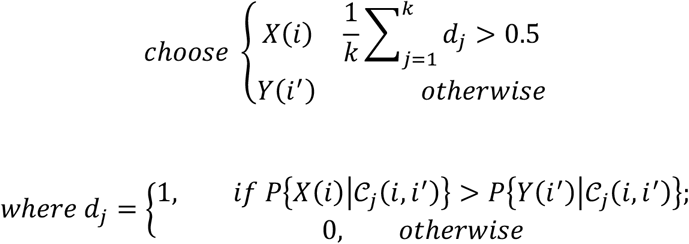

**Equation 2:** Decision Process in Resolving Base-Pair Mismatches of Similar Quality Score

Where *X*(*i*) represents the F sequence with base A of the disagreement, while *Y*(*i*′) represents the R sequence with base B of the disagreement. The element *d*_j_ = 1(0) represents the either the forward (or reverse) sequence base being correct based on Bayesian decision theory, while *C_j_*(*i, i*^′^) represents the window of k-mers considered for both bases.

In the event of a presence of an indel, Levenshtein distance calculations (8) were used by building the final consensus sequence up to the indel point. These then performed the aforementioned comparison methods to decide if the error is either an insertion or deletion, and either skipping or including the base in each respective circumstance.

### DNA sequence translation

In the translation of the consensus DNA sequences into their respective amino acids, sequences still containing ambiguous (N) bases are first removed, and the base array can be reshaped into an n x 3 array to produce a digit corresponding to an entry in a reference amino acid index. The series of digits are then converted to a base 10 number using a dot product with the 3 × 1 vector [4^2^,4^1^,4^0^] with the resulting values compared against a conversion array to generate the peptide sequence (Figure 4 below and Algorithm 5 in Supplementary Information). Regex groups are then also used to determine the presence and location of barcodes and trim accordingly, while simultaneously removing erroneous bases.

**Figure 4.**
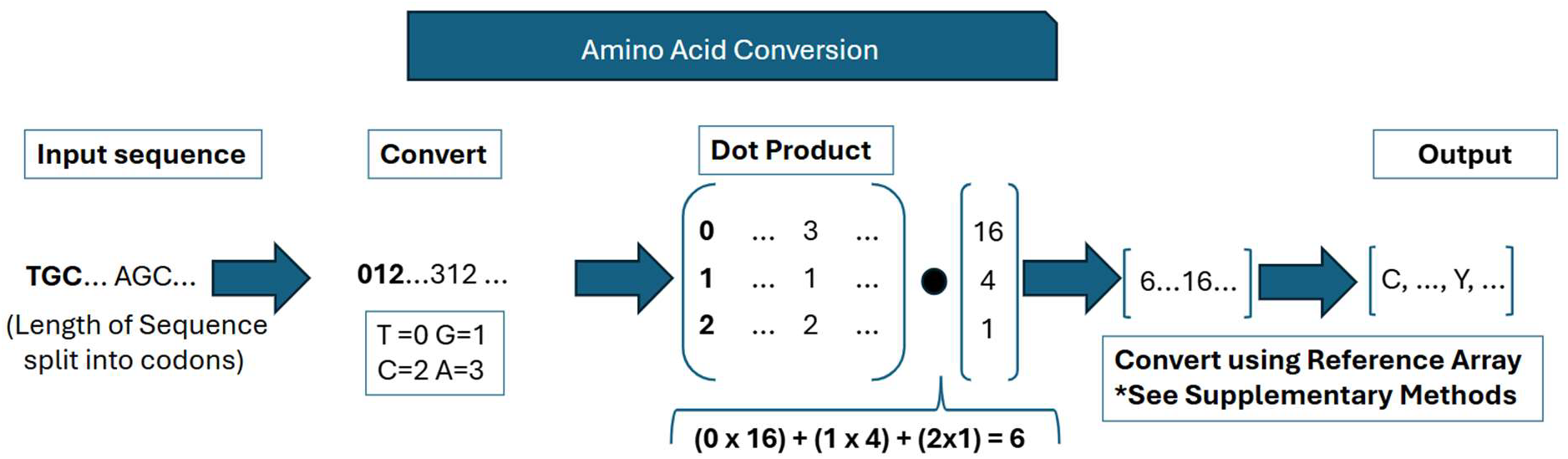
Exemplar use of process filtering and converting codon sequences into amino acid sequences in P3ANUT. Initially, low-quality sequences are removed to optimize processing. Codons, sets of three nucleotide bases, are checked for the presence of ‘N’ (indeterminate bases) using a minimum value check. Sequences with high numbers of low-quality bases are excluded before conversion to reduce data size and enhance runtime efficiency. The sequence encoding allows for base-4 number conversion by subtracting 1 from the array and reshaping it into a matrix. A dot product operation maps each codon to a base-10 index that matches a reference array and then outputs an amino acid sequence.

Before converting the codon sequences into amino acids sequences, low quality sequences need to be removed. Amino acids are determined by a series of 3 bases. A N case within the protein sequence leads to an indeterminate conversion. Through checking the minimum value of the encoded base, the script is quickly able to determine if an N base is present. Sequences that contain many low-quality sequences do not represent the issue with amino acid conversion. Rather, removing them before conversion saves on run time through reduction in the size of data. This is efficiently performed by a Boolean “less than” operation and calculating the number of non-zero or false terms. With the removal of any sequence containing an N base, the sequence base encoding can have 1 subtracted from the array. This allows the interpretations of a codon base as a base 4 number. The sequence base array can be reshaped into an N x 3 array. Each element in the 1 × 3 array can be thought of as a digit in a reference amino index. A dot product can efficiently and effectively convert the series of digits into a base 10 number. The resulting m value of the dot product is used to correlate to the index of a predefined value within a conversion array. To determine the priority of each encoded sequence, optional regex groups are used. A regex statement simultaneously accomplishes two tasks within the same operation. The statement can determine the presence of the barcodes and remove erroneous bases. The presence of the optional groups directly correlates to associated case and priority. The number of these groups directly correlates to the priority.

### Sequence counter

From the resulting dataset of given peptide sequences and their frequencies a sequence counting process can be conducted, as summarised in Figure 5. The sequence counter script provides a modular framework for analysing and counting sequences (DNA or amino acids) extracted from JSON or CSV files. The main functions include **countJsonFile** and **countCSVFile**, which handle file parsing to extract sequences based on specified tags or columns and then pass the sequences to the core **parse** function for processing.

**Figure 5.**
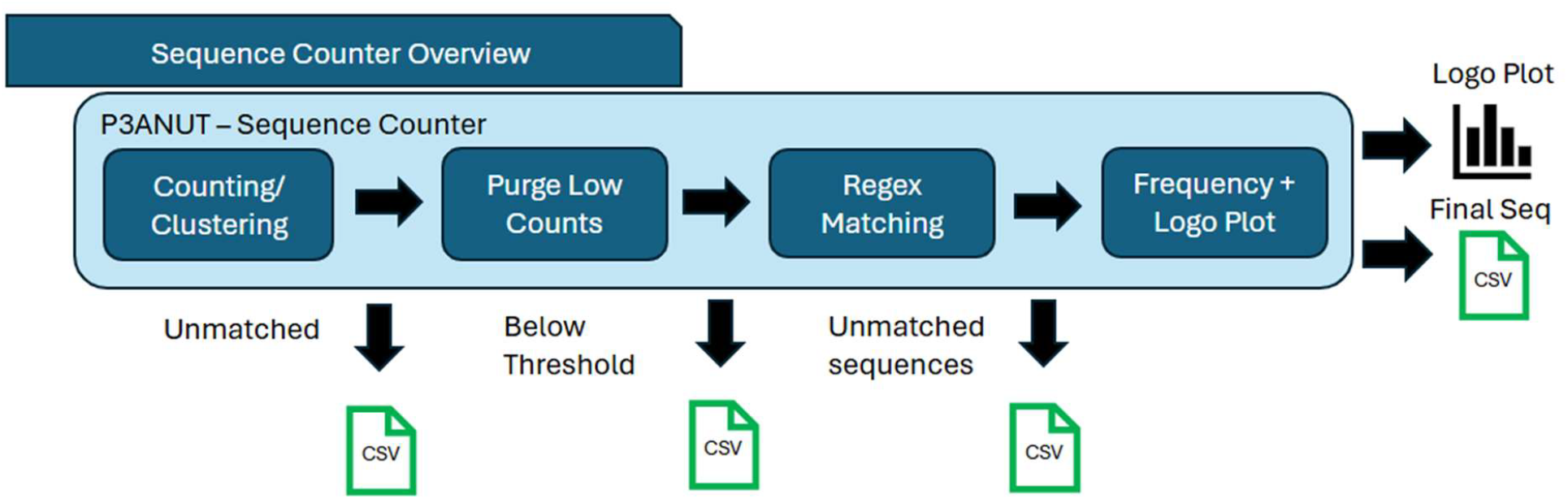
Summary of Sequence Counter Script. The sequencing counting script provides a series of steps to filter out sequences that fail to meet criteria provided by the user, such as sequences without enough hits required, or failure to match a given regular expression, such as a section of the peptide that is always consistent. The outputs are logo plots and final sequence collection files for further processing.

The **parse** function is the centrepiece of the script, orchestrating sequence processing based on user-specified parameters. It supports multiple encoding methods, such as ONEHOT or None, and counting methods, including direct, DBSCAN, and OPTICS. The workflow begins with encoding the sequences into a suitable format, followed by clustering or direct comparison to generate a Data Table of sequence counts. The results are then thresholded to filter out sequences below a specified count, and regex patterns are applied to match sequences of specific lengths.

Output files include purged CSVs for low-count sequences, matched sequences grouped by regex-defined lengths, and optionally sequence logos or tree visualizations for pattern analysis. The script incorporates error handling to manage missing sequence tags or unsupported methods and allows extensive customization via keyword arguments, such as adjusting sequence barcodes, length thresholds, and clustering parameters.

By integrating clustering, regex-based sequence matching, and visualization capabilities, the sequence counter is highly adaptable for bioinformatics applications, enabling flexible and robust analysis of DNA and protein sequence datasets.

### Downstream Analysis

After data processing and translation is completed, P3ANUT offers an optional analytical pipeline that allows for the loading of multiple biological repeats of a given sample. Here, we disclose an example of the selection of cyclic 7-mer mannose glycopeptides against the mannose-binding lectin, ConA using custom-built cyclic 7-mer libraries. Two screening rounds were undertaken in triplicate by panning in parallel three forms of the library (Figure 6a). In Group A, the aim was to identify sequences that bind specifically to the lectin, ConA. It involved screening library 1, which was derivatised with mannose, against ConA and bovine serum albumin, which is not known to bind mannose (i.e. a non-binding control). In Group B, we sought to identify glycopeptides that bind specifically to the carbohydrate recognition domain (CRD). This was achieved by comparing Library 1 with an underivatized library (library 3) which presented only peptides, which may bind to any site on the lectin or possibly in a glycan-independent manner to the accessory binding sites. With Group C we aimed to identify glycan-specific sequences. This involved a comparison of the outputs from Library 1 to the screening outputs of a library derivatised with galactose which presented the non-cognate galactose glycopeptides (Library 2). Identification and exclusion of off-target binding sequences facilitates the stringent selection of glycopeptides that bind specifically to the CRD of ConA and the accessory binding site (shown schematically in Figure 6b).

**Figure 6.**
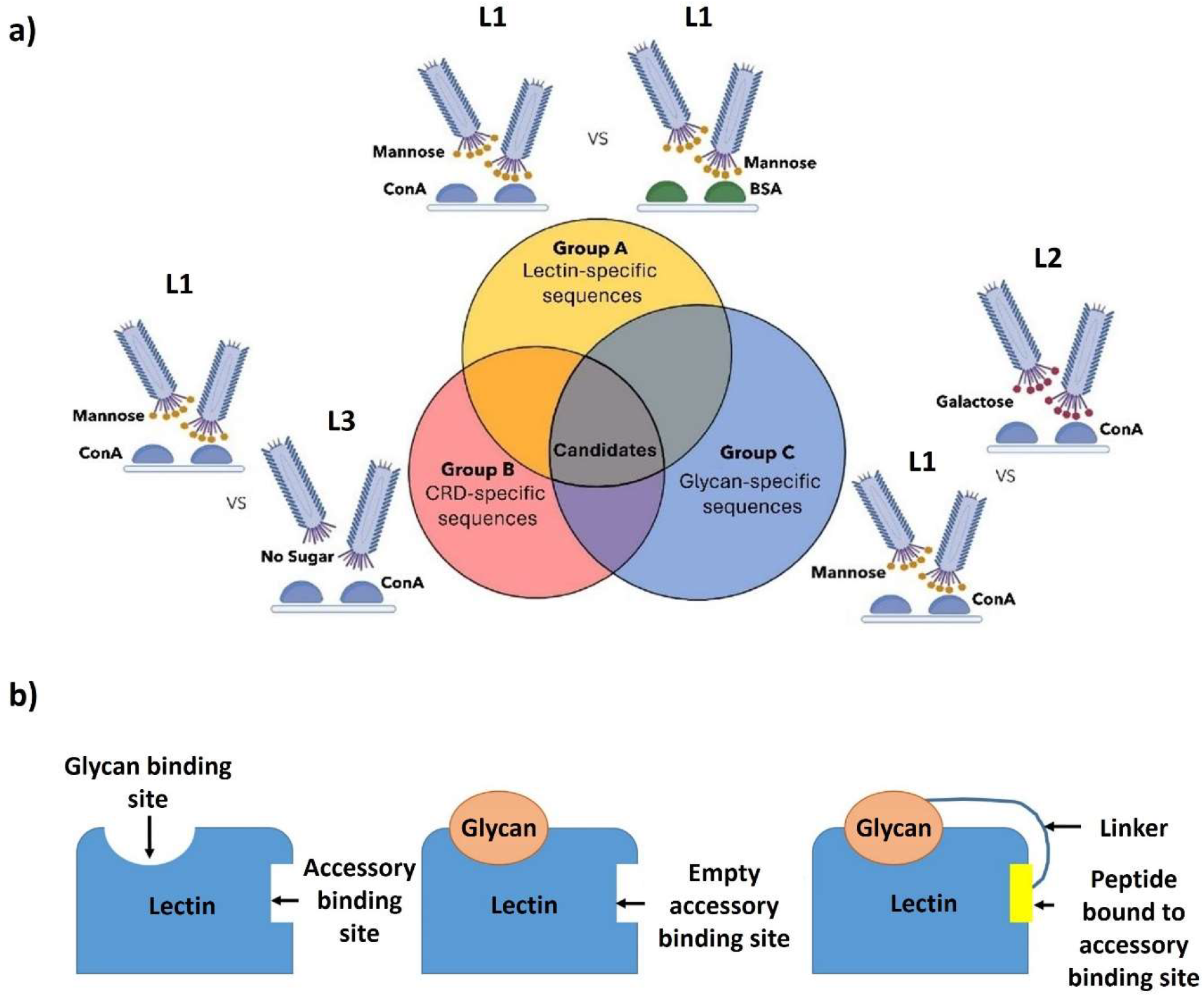
Library screening approach against ConA. (a) Parallel library screens are undertaken using three different library forms – one with the cognate glycan (mannose) appended (L1); one with a control glycan (galactose) appended (L2); and a third with no glycan appended (L3). Comparison of the three pairwise comparisons produces a candidate list for further investigation. (b) Schematic diagram showing the general glycopeptide display approach to identify ligands that occupy both the native glycan binding site (aka carbohydrate recognition domain, CRD) and nearby accessory binding site to increase ligand binding affinity.

The unamplified round 2 eluates were analysed using P3ANUT to yield unique peptide sequence lists that were accompanied by their mean differential abundance ratios and p-values derived from t-tests. These were inputs for the creation of Volcano plots that can be customised by the user to enable facile candidate identification in the quadrant counts.

In our example, we sought to identify only mannose glycopeptides that bind specifically to the carbohydrate recognition domain of ConA though complementary interactions between the cognate glycan fragment and the accessory binding site. Figure 7 shows an example of a pairwise comparison of the outputs from a cognate mannose glycopeptide screen against ConA and the same library screened against the control protein, BSA (i.e. Group C, Figure 6a). Only 682 peptides sequences were identified at a significance level of p<0.05 and at an enrichment level of >5, which is equivalent to 0.231% of the total number of unique sequences detected by Illumina NGS. P3ANUT allows for easy adjustment of the cutoffs of both values depending on the strictness the user desires and production of graphs reflecting such, as well as the export of the peptides found in any of the four quadrants separately for further processing.

**Figure 7.**
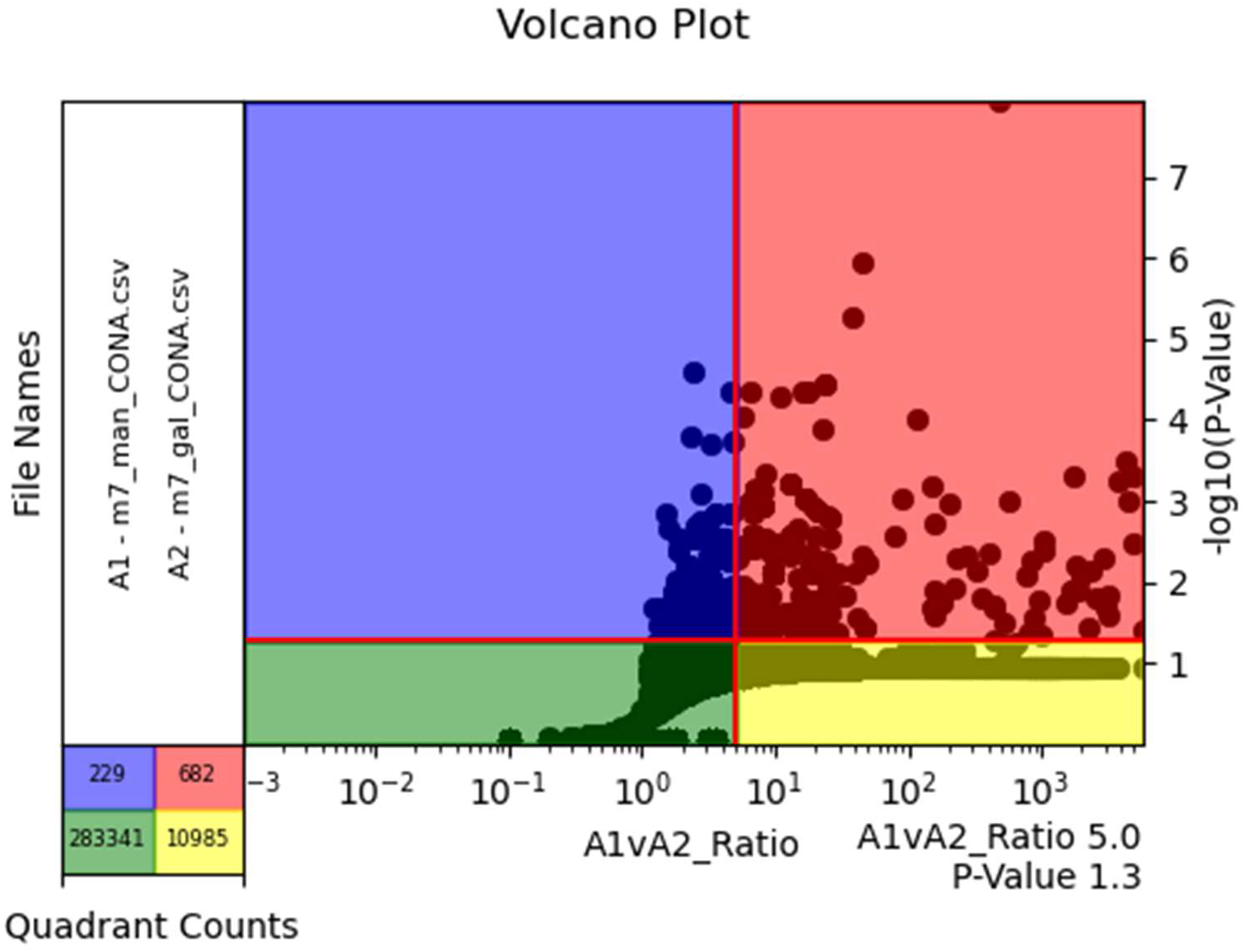
Volcano plot showing the enrichment of mannose functionalised monocyclic 7-mer peptides against ConA versus BSA after two screening rounds (Group C, Figure 6a). Candidates are filtered out from the initial sequencing outputs from Library 1 screening against the ConA target (File A1) and the galactose library 2 against ConA (File A2), achieved by dividing the sequence counts from Files A1 by that in A2. Statistical significance was measured with a one-sided t-test and is represented on the Volcano plot as the –log of the resulting p-value.

We embedded an Upset plot generator that compares the intersections of multiple pairwise comparisons to reveal the binding sequences that are enriched exclusively in the target group and represent candidates for further investigation i.e. 162 unique sequences in *GroupA ∩ GroupB ∩ GroupC* ranked by their abundance (see Figure 8, also Section 5 in Supplementary Information).

**Figure 8.**
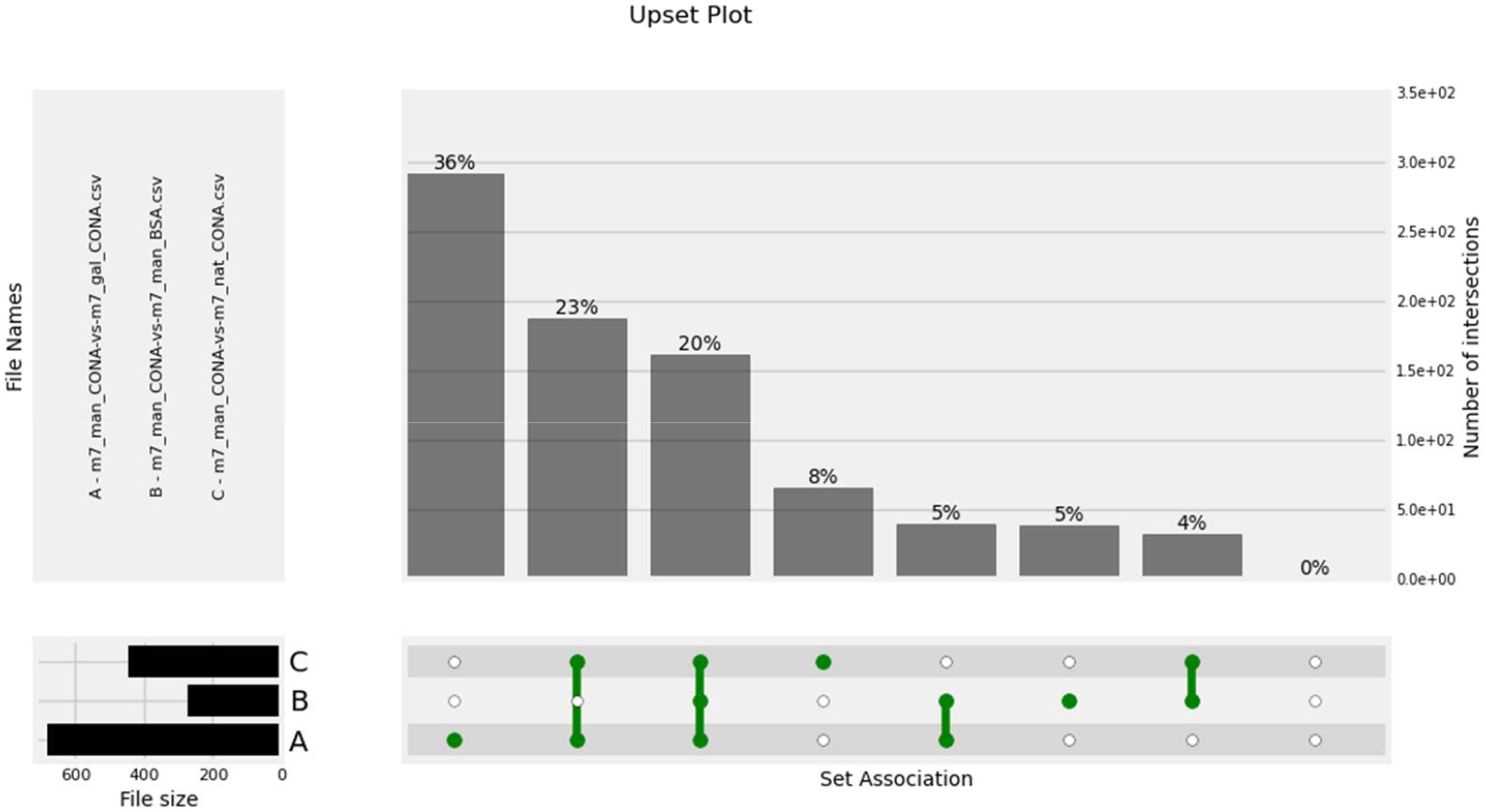
Use of UpSet Plot for identification of candidate sequences. Note that the UpSet plot enables easy dataset comparisons and specific filtering of candidates lists. For Group A,B,C in Figure 6a, their intersection A ∩ B ∩ C comprises 162 unique, candidate mannose glycopeptide sequences by eliminating off-target binding sequences.

### Runtime Improvements

In our stated example, three, pairwise comparisons of sequencing outputs, each present in triplicate, delivered nine sets of forward and reverse sequencing reads. Each consistently contained more than one million sequence pairs, with an average FASTQ file size of 758 MB and equivalent to more than 100x coverage of each clone post-panning. The script runtime was measured by running the single- and multi-processor versions of P3ANUT as well as Step1+2 of the scripts from Rebollo et al. (5) and repeated against each paired dataset three times to find means and standard deviations (Figure 9). Against the compared pipeline, the single-processor version of P3ANUT is at minimum 15.9% slower, with this difference increasing by approximately 0.032% for every additional million sequences in a given file on average. By comparison, the multi-processor version of P3ANUT is at minimum 65.6% faster, though this difference decreases by approximately 0.364% per million sequences, reaching parity with the compared pipeline at around 129 million sequences where on average both pipelines are expected to have a comparable runtime. Given an average file size of approximately 1 GB per sample, a large sequencing run consisting of 70 samples will take slightly over two hours to process fully using the multi-processor function of P3ANUT on average consumer hardware.

**Figure 1.**
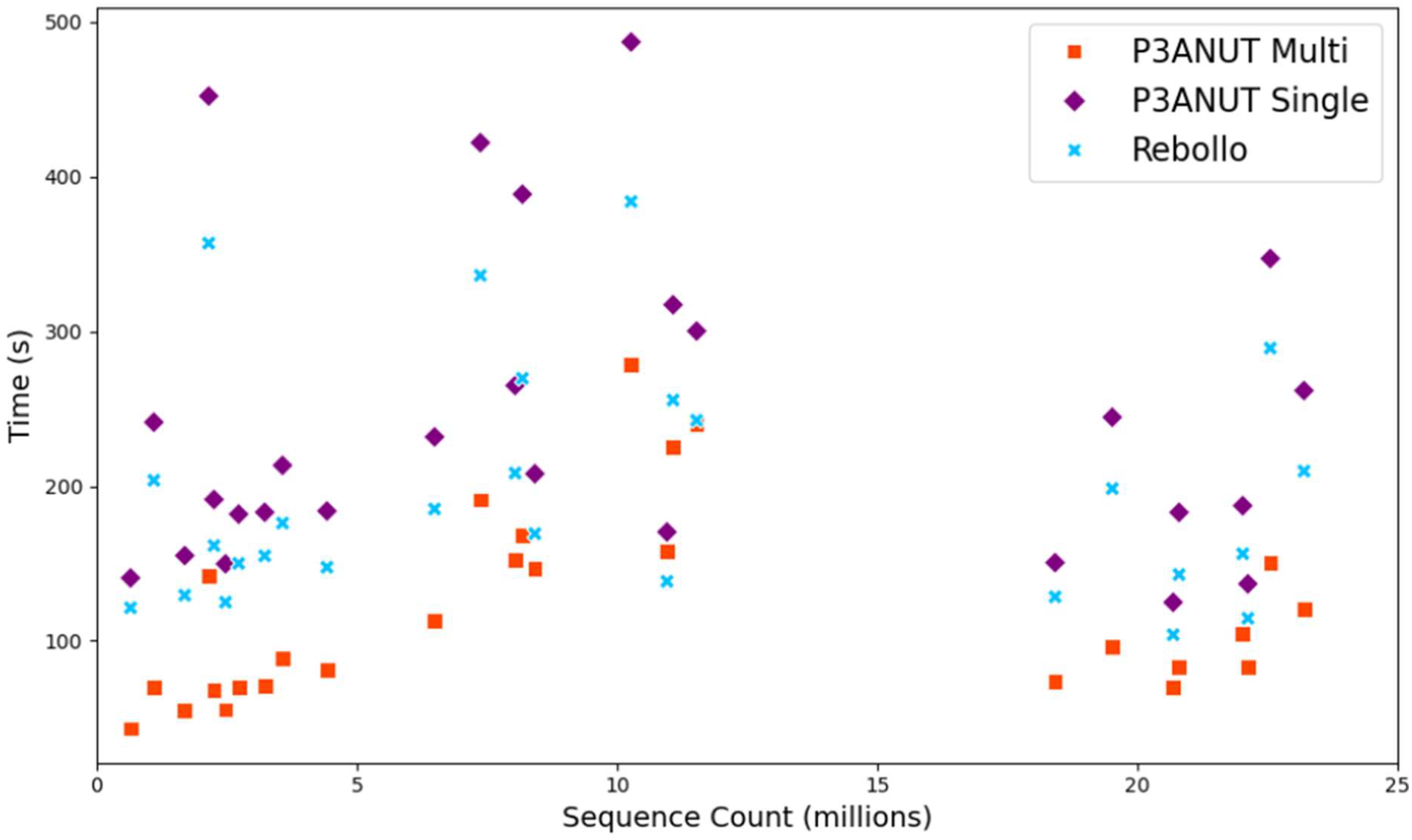
Runtime Performance of P3ANUT against Rebollo Pipelines. Two versions of P3ANUT (the single-and multi-processor versions) are compared against the runtimes of Step1 and 2 from Rebollo et al (5). Using FASTQ files with sequence sizes up to approximately 22 million sequence counts. Note that in addition to sequence sizes, the performance is also subject to other factors, including parameters and the complexities of the dataset.

Figure 9 shows the performance metrics of the single- and multi-processor versions of P3ANUT compared to the combined runtimes of Step1+2 from Rebollo et al (5). All three pipelines show a linear growth (O(n)) in runtime, indicating neither script has functions that cause worse time complexity. For all the tested files, multi-processor version of P3ANUT has the best runtime performance, followed by Rebollo, and then the single-processor version of P3ANUT. Both versions of P3ANUT follow the same trend in runtime as file size increases, which is different from the pipeline of Rebollo et al.

### Pipeline Read Retention and Quality Comparisons

Read retention was also measured across both pipelines by comparing the total sequence numbers across the forward and reverse files. In the case of Rebollo et al. (5), the average of reads retained in both files was used; for P3ANUT, the number of pairs retained was compared to the maximum theoretical pairings. Accounting for this, P3ANUT shows a read recovery rate of approximately 97.5% when including non-pairs that pass quality control (and 96.4% when excluding these), compared to the approximately 65% recovery rate seen in the scripts presented in Rebollo et al. Alongside this, erroneous sequences can be detected if they show incorrect peptide expression in the sections of the peptide that do not change i.e. the fixed N terminal SC- and the C terminal linker sequence, -CGGG. Erroneous sequences are filtered out in most cases by both pipelines using regex matching.

## DISCUSSION

NGS affords a deep analysis of phage display library diversity before and after screening, thus enabling the performance of significantly more complex screening studies with multiple groups. This may be achieved through parallel screening against different targets and controls in multiwell plates or simultaneous screening using silently barcoded libraries as shown by Derda and co-workers (14). By comparing the output from two different panning targets it can be assumed that non-specific binders are removed from the candidate pool.BSA functions as a control for this case, but others can be used such as IgG or, depending on the target, a closely related or mutant protein. In pursuit of glycopeptide binders for ConA our screen was designed such that a tripartite screen was undertaken to exclude: (1) mannose glycopeptides that bound non-specifically to the BSA control; (2) non-glycosylated peptides that bound to sites distal from the CRD, and; (3) non-cognate (galactose) glycopeptides that might bind to distal sites principally through peptide-protein interactions. Notably, it is possible that there may be some sequence overlap between the second and third options listed above because these binding interactions would be glycan-independent.

To enable rapid and accurate interrogation of phage display screening outputs we sought to develop a pipeline that improves the fidelity of the DNA sequencing outputs such as base substitutions, insertions and deletions (indels). These errors have strong potential to diminish sequencing fidelity and therefore have negative impacts on screening studies. Indels and substitutions cause downstream frameshifts that result in fictitious peptide sequences emerging from the screen and wastes time and money that is invested in pursuing irrelevant hits. Traditionally, 3-4 rounds of library screening were undertaken to strongly enrich for strong binding clones such that each binding clone is present in large numbers. In recent times, growing awareness of the issue of parasitic phage clones has prompted a move to shorter screening protocols comprising only one or two rounds to prevent parasitic clone overgrowth and masking of true target binding clones. Yet, when single screening rounds are undertaken, the copy number of each clone is small, which magnifies the deleterious impact of substitutions and indels on peptide hit selection.

The pipeline presented in this work is demonstrated to show both significantly improved read retention and process runtimes when judged against a competitor pipeline for typical phage display datasets. This was achieved using a combination of Levenshtein distance calculations and k-mers to resolve base disagreements. The inclusion of a graphical user interface in addition to a command line interface and readily adjustable analytical scripts is also a first in the field. P3ANUTwill permit even those unfamiliar with bioinformatics to use this software and develop high-quality analysis. The use of innovations first used in other fields for phage display has been seen previously with the utilisation of Primer ID (15) to detect incorrect reads library screens (6). Yet, this approach resulted in an insignificant increase in recovery rates.

Although the multi-processor version of P3ANUT has a favourable runtime for typical phage datasets, it might be slower for large-scale datasets, that is, those with approximately 25 million sequence count and above. In a future work, we will explore how runtime can be further improved by the implementation of GPU processing for large-scale matrices determining both combined read quality as well as resolution of base conflicts. We found that on hardware with low RAM installation or at times when less than 2 GB of RAM is free, P3ANUT faces a risk of out-of-memory errors and run cancelation. In the future this problem could be mitigated by file fractionation and loading them into memory individually rather than as full sequencing files.

Here, we aimed to provide an automated pipeline that is simple to use and delivers easily interpretable results for non-bioinformaticians, built using a free-to-use language that presents opportunity for future development. By integrating robust error-correction through Levenshtein distance calculations and k-mer-based sequence validation, P3ANUT dramatically improves read fidelity and sequence retention rates while maintaining processing efficiency. The ability to correct for sequencing errors, particularly those stemming from indels and substitutions, ensures that researchers can derive more accurate candidate lists for downstream applications, thereby reducing the risk of false positives and improving the overall reliability of phage display library screens.

Accessibility is one of the key strengths of the P3ANUT pipeline. By offering both command-line and UI-based interfaces, the tool accommodates a broad user base, from computational biologists to experimental researchers with minimal bioinformatics experience. Furthermore, its compatibility with multiple display technologies, including mRNA and yeast display, demonstrates its versatility and broad applicability beyond phage display sequencing. This adaptability positions P3ANUT as a powerful tool for researchers engaged in a range of ligand discovery and biomolecular engineering applications.

The implications of this work extend beyond error correction and sequence processing. The ability to filter non-specific binders and identify high-affinity candidates with greater confidence will expedite efficient peptide drug discovery workflows. Despite these advancements, future work could focus on further optimizing computational efficiency. While the multi-processor implementation already enhances speed, rewriting core components in lower-level programming languages such as C or Rust could further improve performance. Additionally, integrating GPU-accelerated processing may enable real-time sequencing analysis, a valuable feature for high-throughput screening applications. Incorporating machine learning techniques to predict high-affinity binders based on sequence motifs could provide another layer of analytical power, enhancing the ability of researchers to prioritize candidate peptides for experimental validation.

In conclusion, P3ANUT addresses a longstanding need for an efficient, user-friendly, and highly accurate tool for peptide display sequencing analysis. By reducing sequencing errors and offering extensive analytical functionalities, it not only enhances existing methodologies but also opens new possibilities for ligand discovery and biomolecular research. As sequencing technologies continue to evolve, P3ANUT provides a strong foundation for future innovations in high-throughput peptide screening and computational biology.

## Supporting information

P3ANUT SI

## DATA AVAILABILITY

The model data used in this work and the P3ANUT pipeline are available online through github.com/taoyangwu/P3ANUT/

The scripts described in this publication are available online through both GitHub (https://github.com/taoyangwu/P3ANUT) and a permanent copy of the version used in this work is provided as part of the Supplementary Data.

## AUTHOR CONTRIBUTIONS

Liam Tucker: Conceptualization, Software, Methodology, Investigation, Supervision, Writing—original draft, review & editing. Ethan Koland: Conceptualization, Software, Methodology, Writing—original draft, review & editing. Jasmyn Gooding: Software, Writing—original draft, review & editing: Hassan Boudjelal: Methodology, Investigation, Resources. Sanaz Ahmadipour: Resources. Robert A Field: Resources, supervision. Derek T Warren: Supervision. David Baker: Investigation, Resources. Yingliang Ma: Resources, supervision. Maria Marin: Supervision, Writing—review & editing. Taoyang Wu: Supervision, Software, Writing—review & editing. Chris Morris: Conceptualization, Supervision, Writing— review & editing.

## FUNDING

LAT was supported by the MRC Doctoral Antimicrobial Research Training iCase Programme (grant number MR/R015937/1). HB was supported by the BBSRC Norwich Research Park Biosciences Doctoral Training Partnership (grant number BB/M011216/1). EK and JG thank the UEA Faculties of Science and Medicine & Health Sciences for funding their PhD studentships.

## CONFLICT OF INTEREST

The authors declare that there are no conflicts of interest involved in the presented work.

## Notes

https://github.com/taoyangwu/P3ANUT

### Competing Interest Statement

The authors have declared no competing interest.

