## Supplementary material for "P3ANUT: An enhanced DNA sequencing analysis platform for uncovering and correcting errors in peptide display library screening": P3ANUT SI

† Joint Authors

Correspondence may also be addressed to: Chris Morris, Tel: +44 (0)20 7753 5853; for Phage Display enquiries and Taoyang Wu, Tel: +44 (0)1603 452954; for computational enquiries.

1. **Library Panning**

Targets are first prepared in concentrations ranging between 10-100 µg $mL^{-1}$ in 0.1 M $NaHCO_3$, pH 8.6 containing required ions at biologically relevant concentrations (1 mM) by adding 150 µL of each solution to each well used per target. The wells were coated by sealing the 96-well plate in a humidified container and incubating overnight at 4 °C with gentle agitation.

Once prepared, the coating solution was poured off from each plate and firmly slapped face down on a paper towel to remove residual solution. Blocking buffer (consisting of filter sterilised pH 8.6 0.1 M $NaHCO_3$, 5 mg.$mL^{-1}$ BSA) was then added to fill each well completely and the plate was incubated for at least 1 hour at 4 °C.

The blocking buffer was poured off and the plate slapped face down on a paper towel to remove residual buffer, the wells were then washed rapidly 6 times with TBST wash buffer (TBS + 0.1% v/v Tween-20) containing metal ions in the above concentrations. After the addition of wash buffer the plate was swirled, then the solution was poured off and the plate slapped face down on a clean paper towel.

A 100-fold representation of the library was diluted in 100 µL TBST and pipetted in each coated well, the plate was then gently rocked at room temperature between 10 and 60 minutes. The solution was poured off and the plate slapped face down on a clean paper towel. Afterwards, each well was washed 10 times with TBST with the solution discarded in the earlier described manner on a clean section of paper towel each time.

The remaining bound phage were eluted from the wells by the addition of a 0.2M glycine-HCl (pH 2.2), 1 mg.$mL^{-1}$ BSA buffer in each well, with the plate gently rocked for 10 minutes. The eluates were pipetted into microfuge tubes and neutralised with 15 µL 1 M Tris-HCl (pH 9.1) and titration checked using 1 µL of eluate, the remaining sample was then either reamplified for additional rounds of panning or put forward for DNA extraction.

2. **Phage DNA Extraction**

To the phage-containing supernatant, 200 µL of 20% w/v PEG/2.5 M NaCl was added, then inverted several times to mix and left to stand for 10-20 minutes at room temperature to precipitate the phage from the solution. The sample was then microfuged at 16,100 *xg* for 10 minutes at 4 °C and the supernatant was discarded, this was then re-spun at 16,100 *xg* again for 2-3 minutes and the remaining supernatant was carefully pipetted away and discarded. The pellet was then suspended in 100 µL of Iodide Buffer (10 mM Tris-HCl (pH 8.0), 1 mM EDTA, 4 M NaI) and mixed through tapping the tube, 250 µL of ethanol was then added and the sample was left to incubate for 10-20 minutes at room temperature to precipitate single-stranded phage DNA.

The sample was then spun in a microfuge at 16,100 *xg* for 10 minutes at 4 °C and the supernatant was discarded. The pellet was then washed with 0.5 mL of 70 %v/v ethanol (stored at -20 °C), re-spun, and the supernatant was then discarded before the sample was then left to dry at room temperature for 10 minutes. DNA was then extracted using a standard phenol/chloroform/isoamyl alcohol extraction and ethanol precipitation, then verified for concentration using a Nanodrop 2000.

#### 3. Illumina Sequencing Preparation

Extracted DNA was then subject to the PCR procedure listed in Table S1 to introduce both a barcode and Illumina adaptors to the DNA region containing the randomised sequence.

**Table S1 | PCR Mixture and Amplification Procedure**

| **Component** | **Volume Required** |
|---|---|
| 5x Platinum SuperFi II buffer | 10 µL |
| 10 mM dNTPs | 1 µL |
| 10 µM L1 Primer | 5 µL |
| 10 µM R1 Primer | 5 µL |
| Platinum SuperFi II DNA polymerase (2 U/µL) | 0.5 µL |
| Phage DNA template (75 ng) | Dependent on original sample concentration |
| $H_2O$ | Add to make up 50 µL total |

**During amplification**

| | |
|---|---|
| Step 1 – 30 Seconds | 98 °C |
| Step 2 – 10 Seconds | 98 °C |
| Step 3 – 20 Seconds | X °C ($T_{anneal}$ of Barcode used from Table S2) |
| Step 4 – 30 Seconds | 72 °C |
| Step 5 – Repeat Steps 2-4, 34 Cycles | |
| Finish – 5 Minutes at 72 °C | |
| Hold at 4 °C | |

L1 and R1 primer (below) include a barcode (Table S2) used are adapted from previous literature (1), including Illumina adapter overhangs that allow for immediate processing in downstream sequencing. As the $T_m$ is only affected by the locus-specific region of the primers, the addition of adapter overhangs does not affect their $T_m$.

Afterwards, DNA was extracted and purified from the PCR mixture through a second round of phenol/chloroform/isoamyl alcohol extraction and ethanol precipitation. DNA concentration was measured using a Nanodrop 2000 and product length was verified using PAGE.

L1 Primer: 5′-TCGTCGGCAGCGTCAGATGTGTATAAGAGACAG-NKKN[BAR]TATTCTCACTCT-3′

R1 Primer: 5′-GTCTCGTGGGCTCGGAGATGTGTATAAGAGACAG-NKKN[BAR]CGAACCTCCACC-3′

**Table S2. Primer Barcodes**

| Primer | Right Primer | | Left Primer | | $T_{anneal}$ (°C) |
|---|---|---|---|---|---|
| | Primer Sequence | $T_m$ (°C) | Primer Sequence | $T_m$ (°C) | |
| BAR1 | NKKN GTA CGA ACC TCC ACC | 55.9 | NKKN GTA TAT TCT CAC TCT | 46.1 | 53.0 |
| BAR2 | NKKN GAC CGA ACC TCC ACC | 58.7 | NKKN GAC TAT TCT CAC TCT | 48.7 | 62.0 |

Once the sequences were confirmed to be of expected length and satisfactory quality and concentration, DNA was processed as a library for sequencing. This was carried out by diluting DNA to 5 ng.µL$^{-1}$. A tagmentation master mix of 0.5 µL Bead-Linked Transposomes (BLT), 0.5 µL Tagmentation Buffer 1 (TB1) and 4 µL $H_2O$, were added per well, and to each well 2 µL of DNA was transferred. The plate was then sealed, centrifuged briefly and then incubated for 15 minutes at 55 °C.

A barcoding PCR master mix consisting of 10 µL Kapa 2G Fast HotStart ReadyMix and 2 µL $H_2O$ per well was made and 12 µL was added to each well. To this, 1 µL of barcode primers (from a 10 µM index stock) then 7 µL of the tagmentation mix (described above) was added. The plate was then resealed, vortex mixed, and spun down once more. Afterwards, the plate contents were amplified with the following PCR profile in Table S3, below:

**Table S3.** Thermocycler conditions for barcoding PCR

| | |
|---|---|
| Step 1 – 3 Minutes | 72 °C |
| Step 2 – 1 Minute | 95 °C |
| Step 3 – 10 Seconds | 95 °C |

| | |
|---|---|
| Step 4 – 20 Seconds | 55 °C |
| Step 4 – 3 Minutes | 72 °C |
| Step 6 – Repeat Steps 3-5, 13 Cycles | |
| Finish – 5 Minutes at 72 °C | |
| Hold at 4 °C | |

Amplified DNA was quality controlled through Qubit using the plate reader of a Quantifluor dsDNA system, then a few samples were selected from the plate to be analysed on an Agilent Tapestation 4200 with a D5000 tape to confirm successful library preparation.

Once library preparation has passed quality control, the pool was made equimolar (based on ng) in an Eppendorf, vortexed, then spun down. The volume was measured and adjusted to 96 µL with $H_2O$. The pooled library was purified and washed using AMPure beads, then concentration and molarity was determined using a Qubit HS assay and a Tapestation D5000 tape, respectively.

From the pooled library a volume was taken to achieve 20 pM. To this volume, 0.5 µL of freshly prepared 2N NaOH was added and the mix made up to 10 µL with Elution Buffer (EB). The mix was then vortexed and pulse spun for 5 seconds. The denaturation mix was incubated at RT for 5 minutes, then 900 µL thawed and chilled Hybridization Buffer (HT1) was added, then the mix was vortexed and pulse spun again for 5 seconds.

To 96 µL of 20 pM library, 1 µL 20 pM PhiX and 1203 µL HT1 was added, then vortexed and pulse spun. The sample was then loaded into the sample loading well of the Illumina cartridge, then sequencing was carried out according to manufacturer protocol.

**4. LOGO Plot Generation of Test Library**

Sequencing of our $(NNK)_3$ library before electroporation revealed a total library diversity of 7939 of 8000 potential amino acid combinations after removal of sequences with erroneous fixed sections. A LOGO plot is generated prior to cleanup (Figure S1) to reveal amino acid distribution, which can be used to indicate amino acids of interest throughout the library sequence.

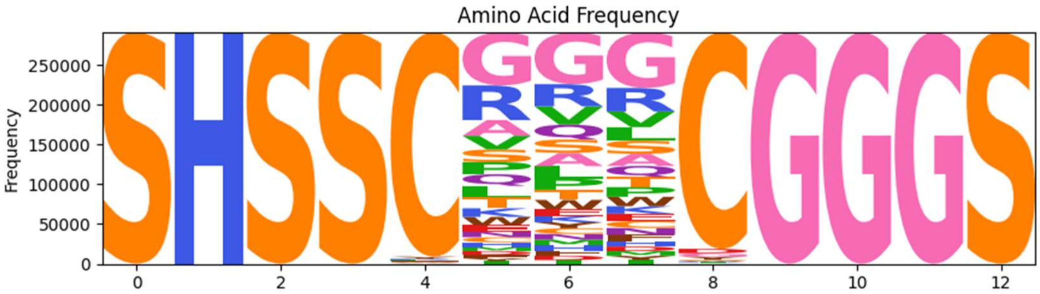


**Figure S1. LOGO Plot of (NNK)$_3$ Library Pre-Electroporation**
Differences in amino acid presence are caused by a combination of number of combinations for each amino acid as well as biases in synthesis of library DNA

### 5. Sequence outputs

Below are the 162 unique, candidate mannose glycopeptide sequences identified by P3ANUT (see Figure 8 in the main text), ranked by their relative abundance (indicated by the mean column).

| sequence | mean | std |
|---|---|---|
| SHSSCGWGGGGWCGGGS | 0.016820988 | 0 |
| SHSSCLRHMTSSCGGGS | 0.004426868 | 0 |
| SHSSCPEHPGPSCGGGS | 0.004401851 | 0 |
| SHSSCKQTAPTGCGGGS | 0.003708842 | 0 |
| SHSSCRQPHQPPCGGGS | 0.00331333 | 0 |
| SHSSCTGQTSARCGGGS | 0.002914168 | 0 |
| SHSSCLFGVTTSCGGGS | 0.002890656 | 0 |
| SHSSCQTSGHTYCGGGS | 0.002768981 | 0 |
| SHSSCQLADPPHCGGGS | 0.002714107 | 0 |
| SHSSCRLPGGNACGGGS | 0.002543704 | 0 |
| SHSSCQTTTQTTCGGGS | 0.002475398 | 0 |
| SHSSCEAPAETSCGGGS | 0.002393584 | 0 |
| SHSSCHKPTPTNCGGGS | 0.002349999 | 0 |
| SHSSCRLPLTNACGGGS | 0.002242903 | 0 |
| SHSSCGGVEGGACGGGS | 0.002028635 | 0 |

| | | |
|---|---|---|
| SHSSCTPPTPTHCGGGS | 0.002005464 | 0 |
| SHSSCSPPSPSTCGGGS | 0.001900799 | 0 |
| SHSSCGEGYVGACGGGS | 0.001749832 | 0 |
| SHSSCWWGGGGACGGGS | 0.001575539 | 0 |
| SHSSCVGVGVVVCGGGS | 0.001561078 | 0 |
| SHSSCKQPTHPTCGGGS | 0.001493224 | 0 |
| SHSSCDKEPKKYCGGGS | 0.001490252 | 0 |
| SHSSCGVREGAWCGGGS | 0.001348778 | 0 |
| SHSSCGGTGIRGCGGGS | 0.001316537 | 0 |
| SHSSCPEIEVWGCGGGS | 0.001266965 | 0 |
| SHSSCTHPPWQTCGGGS | 0.00122045 | 0 |
| SHSSCDGWWAGVCGGGS | 0.001122144 | 0 |
| SHSSCGGGGDRWCGGGS | 0.001106422 | 0 |
| SHSSCGAGGGSGCGGGS | 0.00105379 | 0 |
| SHSSCGDSRGGDCGGGS | 0.00094853 | 0 |
| SHSSCGGGGLWGCGGGS | 0.000810456 | 0 |
| SHSSCGGVWGGECGGGS | 0.000762544 | 0 |
| SHSSCGGLGGKECGGGS | 0.000742896 | 0 |
| SHSSCGGGDGGRCGGGS | 0.000729454 | 0 |
| SHSSCEGEGWGRCGGGS | 0.000722727 | 0 |
| SHSSCGGGGGGRCGGGS | 0.000715976 | 0 |
| SHSSCEGRGGGGCGGGS | 0.000627992 | 0 |
| SHSSCVWGGGVGCGGGS | 0.00062681 | 0 |
| SHSSCGGGVWGDCGGGS | 0.000609885 | 0 |
| SHSSCGEQRFGWCGGGS | 0.000601711 | 0 |
| SHSSCGGERGGKCGGGS | 0.000590286 | 0 |
| SHSSCGGVVGGRCGGGS | 0.000579745 | 0 |
| SHSSCGGGGARGCGGGS | 0.000485262 | 0 |
| SHSSCGGWGGGRCGGGS | 0.000442385 | 0 |
| SHSSCGGGSAGGCGGGS | 0.000333833 | 0 |
| SHSSCGGVRGGGCGGGS | 0.000321349 | 0 |
| SHSSCGGGSGGGCGGGS | 0.00030641 | 0 |
| SHSSCGGGGVGGCGGGS | 0.000220919 | 0 |
| SHSSCAGGGGGGCGGGS | 0.000181796 | 0 |
| SHSSCGGVGGGGCGGGS | 0.000153399 | 0 |
| SHSSCGAGGGRWCGGGS | 0.000111026 | 0 |
| SHSSCGRGGGGWCGGGS | 9.45E-05 | 0 |
| SHSSCGWGGGGRCGGGS | 4.01E-05 | 0 |
| SHSSCGCGGGGWCGGGS | 3.57E-05 | 0 |
| SHSSCGWCGGGWCGGGS | 3.44E-05 | 0 |
| SHSSCGWGGGCWCGGGS | 3.42E-05 | 0 |
| SHSSCGWGGWGWCGGGS | 2.88E-05 | 0 |
| SHSSCWWGGGGWCGGGS | 2.86E-05 | 0 |
| SHSSCPHSPHPLCGGGS | 2.59E-05 | 0 |
| SHSSCGKGLIAQCGGGS | 2.58E-05 | 0 |
| SHSSCEVPWMKRCGGGS | 2.58E-05 | 0 |
| SHSSCGGGRGVGCGGGS | 2.36E-05 | 0 |

| | | |
|---|---|---|
| SHSSCGWGRGGWCGGGS | 2.30E-05 | 0 |
| SHSSCLFCVTTSCGGGS | 2.22E-05 | 0 |
| SHSSCPEHPCPSCGGGS | 2.14E-05 | 0 |
| SHSSCKQPATPPCGGGS | 2.12E-05 | 0 |
| SHSSCQLPATPPCGGGS | 2.10E-05 | 0 |
| SHSSCQQPATPTCGGGS | 2.06E-05 | 0 |
| SHSSCGKGLNSQCGGGS | 2.04E-05 | 0 |
| SHSSCTHPSTSTCGGGS | 1.99E-05 | 0 |
| SHSSCGSGGGGRCGGGS | 1.90E-05 | 0 |
| SHSSCLRHMNSSCGGGS | 1.74E-05 | 0 |
| SHSSCQTSCHTYCGGGS | 1.74E-05 | 0 |
| SHSSCGGTGFRGCGGGS | 1.71E-05 | 0 |
| SHSSCGWGGGGCCGGGS | 1.62E-05 | 0 |
| SHSSCNTTSDKNCGGGS | 1.59E-05 | 0 |
| SHSSCSGQTSARCGGGS | 1.58E-05 | 0 |
| SHSSCWGGAEGRCGGGS | 1.56E-05 | 0 |
| SHSSCRLAGGNACGGGS | 1.49E-05 | 0 |
| SHSSCTPPTPTLCGGGS | 1.47E-05 | 0 |
| SHSSCGGRGIRGCGGGS | 1.46E-05 | 0 |
| SHSSCEVPLKKRCGGGS | 1.43E-05 | 0 |
| SHSSCEELAPQTCGGGS | 1.43E-05 | 0 |
| SHSSCGVRVGAWCGGGS | 1.43E-05 | 0 |
| SHSSCMRRHTNNCGGGS | 1.38E-05 | 0 |
| SHSSCRMPLTNACGGGS | 1.37E-05 | 0 |
| SHSSCVGVLGRACGGGS | 1.37E-05 | 0 |
| SHSSCQTSGHSYCGGGS | 1.35E-05 | 0 |
| SHSSCVGVMKTICGGGS | 1.34E-05 | 0 |
| SHSSCEGVGVRGCGGGS | 1.32E-05 | 0 |
| SHSSCKQSTHHTCGGGS | 1.32E-05 | 0 |
| SHSSCKQTAPTCCGGGS | 1.30E-05 | 0 |
| SHSSCQQGACARCGGGS | 1.30E-05 | 0 |
| SHSSCDKETKKYCGGGS | 1.28E-05 | 0 |
| SHSSCLLPGGNACGGGS | 1.28E-05 | 0 |
| SHSSCKQTAQTGCGGGS | 1.28E-05 | 0 |
| SHSSCNQTAPTGCGGGS | 1.28E-05 | 0 |
| SHSSCQLGASARCGGGS | 1.28E-05 | 0 |
| SHSSCLLHMTSSCGGGS | 1.25E-05 | 0 |
| SHSSCKNPTHTTCGGGS | 1.24E-05 | 0 |
| SHSSCRLPGGIACGGGS | 1.24E-05 | 0 |
| SHSSCMGNNASWCGGGS | 1.23E-05 | 0 |
| SHSSCGEGFVGACGGGS | 1.21E-05 | 0 |
| SHSSCTLPDGQTCGGGS | 1.21E-05 | 0 |
| SHSSCHKTTPTNCGGGS | 1.21E-05 | 0 |
| SHSSCLFVVTTSCGGGS | 1.21E-05 | 0 |
| SHSSCLRHLTSSCGGGS | 1.18E-05 | 0 |
| SHSSCQPTTPQTCGGGS | 1.10E-05 | 0 |
| SHSSCEATAETSCGGGS | 1.06E-05 | 0 |

| | | |
|---|---|---|
| SHSSCTEHPGPSCGGGS | 1.06E-05 | 0 |
| SHSSCPEHPGRSCGGGS | 1.06E-05 | 0 |
| SHSSCQAPAETSCGGGS | 1.06E-05 | 0 |
| SHSSCGGTGISGCGGGS | 1.05E-05 | 0 |
| SHSSCQQPETPPCGGGS | 1.04E-05 | 0 |
| SHSSCTHRSTATCGGGS | 1.04E-05 | 0 |
| SHSSCTPLTQHLCGGGS | 1.04E-05 | 0 |
| SHSSCGGVGRVECGGGS | 1.03E-05 | 0 |
| SHSSCTSPPHPTCGGGS | 1.02E-05 | 0 |
| SHSSCQHPATPPCGGGS | 1.02E-05 | 0 |
| SHSSCHKQTPTNCGGGS | 9.95E-06 | 0 |
| SHSSCGWAGGGWCGGGS | 9.95E-06 | 0 |
| SHSSCGEGGGSGCGGGS | 9.95E-06 | 0 |
| SHSSCQTPKWPFCGGGS | 9.95E-06 | 0 |
| SHSSCPEIEVWVCGGGS | 9.95E-06 | 0 |
| SHSSCDNEPKKYCGGGS | 9.95E-06 | 0 |
| SHSSCWLGGGGACGGGS | 9.95E-06 | 0 |
| SHSSCQSPASPWCGGGS | 9.95E-06 | 0 |
| SHSSCKAPAETSCGGGS | 9.95E-06 | 0 |
| SHSSCGDGYVGACGGGS | 9.95E-06 | 0 |
| SHSSCGGGGLGGCGGGS | 9.83E-06 | 0 |
| SHSSCTRTEAGFCGGGS | 9.78E-06 | 0 |
| SHSSCQQGESARCGGGS | 9.74E-06 | 0 |
| SHSSCPLPSPRGCGGGS | 9.74E-06 | 0 |
| SHSSCKQTAPSGCGGGS | 9.74E-06 | 0 |
| SHSSCRQPHQQPCGGGS | 9.31E-06 | 0 |
| SHSSCGQGYVGACGGGS | 9.31E-06 | 0 |
| SHSSCGGTGIWGCGGGS | 9.31E-06 | 0 |
| SHSSCLRNMTSSCGGGS | 9.31E-06 | 0 |
| SHSSCRVPGGNACGGGS | 9.31E-06 | 0 |
| SHSSCSPPSQSTCGGGS | 9.31E-06 | 0 |
| SHSSCGDQRGGDCGGGS | 9.31E-06 | 0 |
| SHSSCTHPPRQTCGGGS | 9.31E-06 | 0 |
| SHSSCKLTAPTGCGGGS | 9.31E-06 | 0 |
| SHSSCTPLTLDLCGGGS | 9.31E-06 | 0 |
| SHSSCQTSGLTYCGGGS | 9.31E-06 | 0 |
| SHSSCNTTTDKYCGGGS | 9.11E-06 | 0 |
| SHSSCEALDPQTCGGGS | 8.95E-06 | 0 |
| SHSSCDGRIKSTCGGGS | 8.48E-06 | 0 |
| SHSSCQQPANPPCGGGS | 8.47E-06 | 0 |
| SHSSCKPTPPPSCGGGS | 8.44E-06 | 0 |
| SHSSCSDVVGRGCGGGS | 8.44E-06 | 0 |
| SHSSCNPPTYPTCGGGS | 8.44E-06 | 0 |
| SHSSCTQQPTRECGGGS | 8.31E-06 | 0 |
| SHSSCSQPHQPPCGGGS | 8.21E-06 | 0 |
| SHSSCKQGASARCGGGS | 8.21E-06 | 0 |
| SHSSCRQPLQPPCGGGS | 8.21E-06 | 0 |

| | | |
|---|---|---|
| SHSSCQTSKIPTCGGGS | 7.97E-06 | 0 |
| SHSSCKWPHPTECGGGS | 7.97E-06 | 0 |
| SHSSCKQTSPTGCGGGS | 7.57E-06 | 0 |
| SHSSCPQPATPPCGGGS | 7.57E-06 | 0 |
| SHSSCVGGCGGWCGGGS | 7.57E-06 | 0 |
| SHSSCKQTAPKGCGGGS | 7.57E-06 | 0 |

### 6. Pseudo Codes

**Algorithm 1** Sequence Encoding Function

```
procedure ENCODING FUNCTION(FileName)
    sequences := READFILE(FileName)
    encodedSeq = []
    for i ← 1, length(sequences) do
        encodedSeq := []
        for j ← 1, length(sequences[i]) do
            seqBases, seqScores := sequences[i][j]
            intBaseRepresentation := CONVERTTOINT(seqBases)
            qScore := CONVERTTOQSCORE(seqScores)
            encodedQScore := (qScore − 33) × 5
            encodedSeq[j] := encodedQScore + intBaseRepresentation
        end for
        encodedSequences[i] := encodedSeq
    end for
    return encodedSequences
end procedure
```

**Algorithm 2** Forward and Reverse Sequence Merge function

```
procedure MERGE(sequence, k, scoreOffset)
    forwardSequence, reverseSequence := sequence
    mergedSequence = []
    if length(forwardSequence) ≠ length(reverseSequence) then
        return MERGEMISMATCH(sequence)
    end if
    forwardScroes := forwardSequence  mod 5
    forwardBases := forwardSequence/5
    reverseScroes := reverseSequence  mod 5
    reverseBases := freverseSequence/5
    for i ← 1, length(forwardSequence) do
        if forwardBase(i) == reverseBases(i) then
            mergedSequence(i) := (reverseScore(i) + forwardScore(i)) × 5 +
forwardBase(i)
        else if forwardScore(i) − reverseScore(i) > scoreOffset then
            mergedSequence(i) := forwardSequence(i)
        else if reverseScore(i) − forwardScore(i) > scoreOffset then
            mergedSequence(i) := reverseSequence(i)
        else
            mergedSequence(i) := KMERS(i, k, forwardSequence, reverseSequence)
        end if
    end for
    return mergedSequence
end procedure
```

**Algorithm 3** Determining Most likely based through Kmers comparison

```
procedure KMERS(i, k, forwardSequence, reverseSequence)
    minKmerIndex := MAXIMUM(0, i − k)
    maxKmerIndex := MINIMUM(length(forwardSequence), i + k)
    kMersTally := 0
    for j ← minKmerIndex, maxKmerIndex do
        if forwardSequence(j) > reverseSequence(j) then
            kMersTally := kMersTally + 1
        else if forwardSequence(j) < reverseSequence(j) then
            kMersTally := kMersTally − 1
        end if
    end for
    if kMersTally > 0 then
        return forwardSequence(i)
    else if kMersTally < 0 then
        return reverseSequence(i)
    else if SUM(forwardSequence/5) < SUM(reverseSequence/5) then
        return reverseSequence(i)
    else
        return forwardSequence(i)
    end if
end procedure
```

**Algorithm 4** Correcting Indels or Sequence length mismatches with Levenshtein

```
procedure MERGEMISMATCH(forwardSequence, reverseSequence, desiredLength)
    operations := LEVENSTEIN(forwardSequence, reverseSequence)
    maxSequence, maxScore := 0
    if Case1 ∨ Case3 ∨ Case5 then
        for deletion, index ← operations do
            newSequence := reverseSequence[0 : i] + reverseSequence[i + 1 : LENGTH(reverseSequence)]
            score, newMerged := MERGE(forwardSequence, newSequence)
            if Score > newScore then
                maxSequence, maxScore := newMerged, score
            end if
        end for
    else if Case2 ∨ Case4 ∨ Case5 then
        for insertion, index ← operations do
            newSequence := reverseSequence[0 : i] + ”N” + reverseSequence[i : LENGTH(reverseSequence)]
            score, newMerged := MERGE(forwardSequence, newSequence)
            if Score > newScore then
                maxSequence, maxScore := newMerged, score
            end if
        end for
    end if
    return maxSequence
end procedure
```

**Algorithm 5** Decoding and Conversion of Codon Sequences into Amino Sequences.

```
procedure FINALIZE(sequence, startBarcode, endBarcode)
    refernceArray := [...]
                        ▷ Preset array of dna sequence and coresponding amino acid
    bestSequence = ""
    for i ← 1,3 do
        tempSequence := []
        sequenceEndingOffset := LENGTH(sequence) − (LENGTH(sequence) − i)
mod 3
        for j ← RANGE(i, sequenceEndingOffset, 3) do
            dnaBlock = sequence[j, j+3]
            aminoIndex = dnaBlock · [16,4,1]^T
            tempSequence(i) := refernceArray(aminoIndex)
        end for
        if startBarcode ∈ tempSequence & startBarcode ∈ tempSequence then
            return tempSequence
        else if sum(tempSequence) > sum(bestSequence) then
            bestSequence := tempSequence
        end if
    end for
    returnbestSequence
end procedure
```
